# Cell cycle repression and DNA repair defects follow constricted migration

**DOI:** 10.1101/240838

**Authors:** Charlotte R. Pfeifer, Yuntao Xia, Kuangzheng Zhu, Dazhen Liu, Jerome Irianto, Shane Harding, Roger A. Greenberg, Dennis E. Discher

## Abstract

Cancer cell invasion into tissue or narrow capillaries often elongates the nucleus and sometimes damages it, but cell cycle effects are unknown and highly relevant to tumorigenesis. Here, nuclear rupture and DNA breaks caused by constricted migration are quantified in different phases of cell cycle - which is effectively repressed. Cancer lines with varying levels of contact inhibition and lamina proteins exhibit diverse frequencies of nuclear lamina rupture after migration, with pre-rupture dilation of gene-edited RFP-Lamin-B1 preceding DNA repair factor leakage in pressure-controlled distension. Post-migration rupture indeed associates with mis-localized DNA repair factors and increased DNA breaks as quantified by pan-nucleoplasmic foci of γH2AX, with foci counts always suppressed in late cell cycle. When contact-inhibited cells migrate through large pores into sparse microenvironments, cells re-enter cell cycle consistent with release from contact inhibition. In contrast, constricting pores effectively delay re-entry, but the excess DNA damage nonetheless exceeds any cell cycle dependence. Partial depletion of topoisomerase does not strongly affect cell cycle or the excess DNA damage, consistent with weak dependencies on replication stress. Constricted migration thus impacts cell cycle as well as DNA damage.

## Introduction

In recent years, there has been a major push to sequence tumor genomes and catalogue the somatic mutations that underlie cancer development [1–5] for use in therapies including immunotherapies. How these somatic mutations arise remains an active topic of study. Vogelstein and coworkers analyzed oncogenic mutation risk in 31 diverse tissues (brain, bone, marrow, lung, etc.), and they proposed that risk increases with DNA replication rate in the resident adult stem cells that normally divide within each normal tissue [6]. However, the correlation is weak (*R*^2^=0.6), which suggests additional mechanisms beyond errors in replication also contribute to cancerous mutations [7]. Genomic variation has been reported to emerge after migration of cancer cells through constricting pores [8] as occurs *in vivo* during invasion and metastasis (Fig. 1Ai,ii).

**Figure 1.**
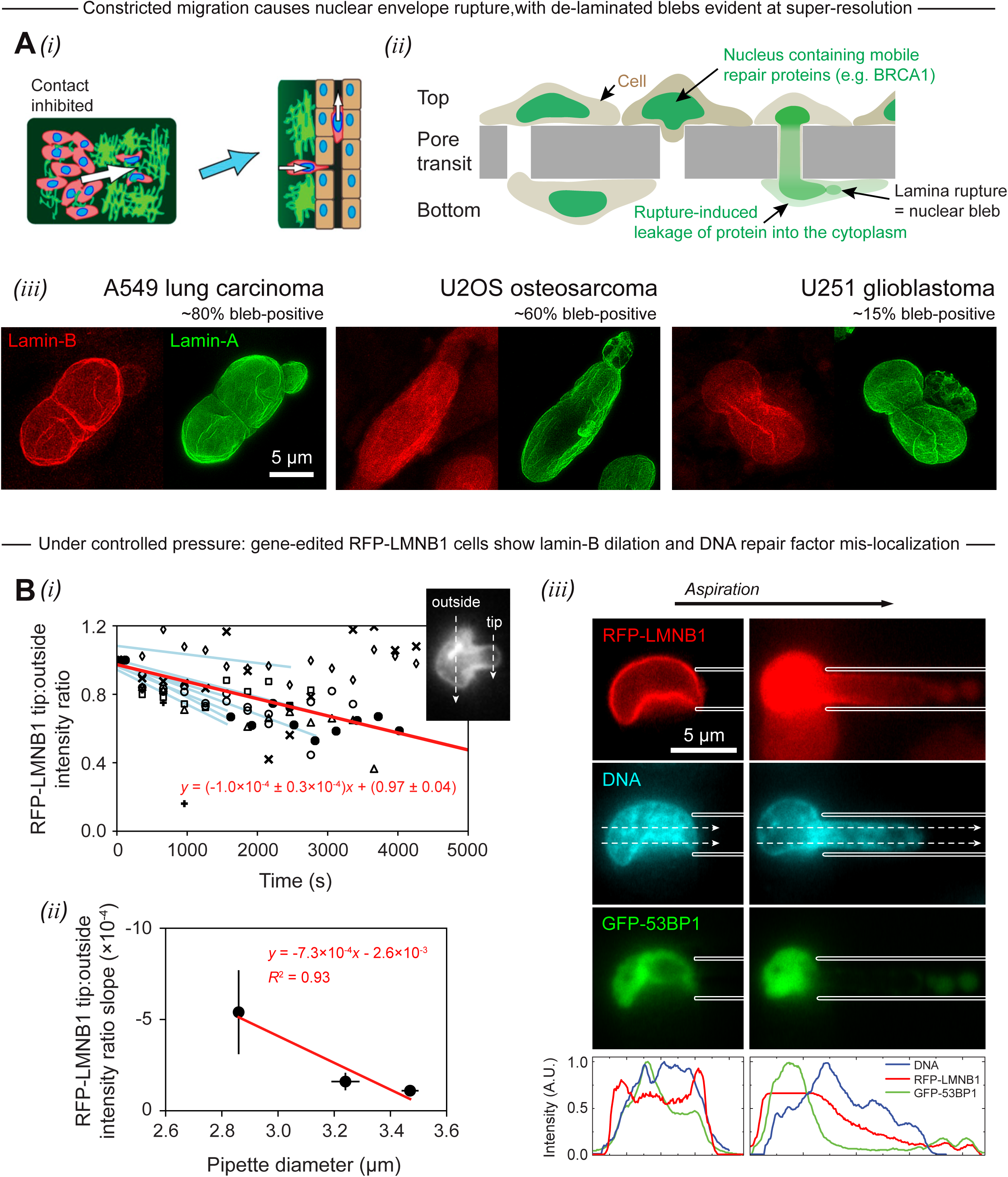
Constricted migration causes dilation of the nuclear lamina, followed by lamina rupture and mis-localization of DNA repair factors. (A) Invasion is a hallmark of cancer, and cancer cells *in vivo* must squeeze into tight spaces as they invade stiff tissue matrix, penetrate basement membrane barriers, and enter distant capillary beds during metastasis. (i) Schematic showing metastasizing cancer cells migrating from a region of high tumor cell density (i.e. the tumor mass) to regions of low tumor cell density, with possible effects on contact inhibition and cell cycle. (ii) Transwell membranes perforated with cylindrical holes of various sizes are used to study cancer cell migration *in vitro*. Cells are seeded on top of the membrane at very high density; driven by over-crowding, the cells migrate to the bottom (a low-density microenvironment) over many hours. During migration through constricting pores, cells exhibit frequent lamina rupture at sites of nuclear blebs, followed by mis-localization of mobile nuclear proteins, including crucial DNA repair proteins, into the cytoplasm. (iii) Super-resolution images show lamina dilation in the nuclear blebs of three different cancer cell lines that have migrated through 3++ μm pores. Percent ‘bleb-positive’ refers to the proportion of cells (of that cell type) that exhibit blebs following 3 μm pore migration. (B) (i) Seven A549 cells with endogenously RFP-tagged LMNB1 were pulled into a ~3 μm pipette under controlled pressure. In all cells, lamin-B shows initial depletion from the leading tip of the nucleus, which is quantified as a decline in the tip-to-outside RFP-LMNB1 intensity ratio (inset) over an ~hour-long aspiration experiment. Each cell is represented by a different symbol and fit with a blue line. The red line is fit to all data. (ii) The blue-line slopes from panel (i) are binned and plotted against pipette diameter (which varies slightly between aspiration experiments). The rate of lamin-B depletion decreases with pipette diameter, suggesting that higher curvature—as imposed on nuclei by smaller pores—causes more rapid dilation of the lamina. (iii) A representative A549 RFP-LMNB1 cell with overexpressed GFP-53BP1 squeezes into a micropipette. The aspirated nucleus shows lamin-B dilation at the leading tip and rupture of GFP-53BP1 into the cytoplasm. Notably, the nucleus also shows segregation of GFP-53BP1, a mobile protein, away from the region of highest chromatin compaction at the pipette entrance.

Migration of various cancer cell lines, immortalized epithelial cells, and some primary cells through narrow channels or small circular pores has been shown to rupture the nuclear envelope, thereby causing cytoplasmic mis-localization of GFP-NLS [9,10] and nuclear factors such as DNA repair factors [8]. Constrictions also cause intra-nuclear exclusion of mobile nuclear proteins (such as DNA repair factors) away from strongly compacted chromatin [11]. For some repair factors, inactivating mutations are such well-established risk factors for cancer that they warrant surgical removal of ovary and breasts [12], and mouse knockouts or heterozygous mutants for repair proteins have also been shown to alter chromosome copy number (e.g. [13]). Constricted migration indeed seems to increase DNA damage based on multiple measures [8]: increased foci of endogenous γH2AX, increased foci of the upstream kinase phospho-ATM, and longer electrophoretic comets consistent with cleaved DNA. However, cell cycle phases are linked to DNA damage, with cell cycle checkpoints ensuring that DNA damage tends to decrease through repair. Such processes might relate somehow to Vogelstein and coworkers’ correlation of replication with cancer risk, and certainly motivate a careful account of both cell cycle and DNA damage after constricted migration.

## Results

After migration through 3 µm pores, a fraction of nuclei (as many as 80% for an A549 lung carcinoma line) exhibit blebs in the nuclear lamina that have been associated with nuclear rupture [9,10]. Super-resolution microscopy of migration-induced blebs shows near complete loss of lamin-B, which confers solidity or elasticity to nuclei [14], as well as dilated webs of lamin-A (Fig. 1A-iii). Micropipettes have a similar geometry as pores and allow controlled pressures to be applied to detached cells, and A549 cells with gene-edited RFP-LMNB1 confirm depletion of lamin-B from the protruding tip of the nucleus over hour-long timescales (Fig. 1B-i). The rate of lamin-B depletion also decreases with pipette diameter (Fig. 1B-ii), which indicates that higher curvature imposed by smaller pores causes greater dilation of the lamina that should favor nuclear rupture. Leakage of normally nucleoplasmic factors is indeed evident with loss of the DNA repair factor GFP-53BP1 from the nuclei of the A549 cells (Fig. 1B-iii).

After constricted migration, mis-localization of endogenous DNA repair factors, such as KU80, is also evident (Fig. 2A-i). Repair factor loss is accompanied by an increase in pan-nucleoplasmic foci of γH2AX and 53BP1, which tend to co-localize in super-resolution images of a pore-migrated U251 glioblastoma cell (Fig. 2A-ii). Indeed, while the mesenchymal-like U251 cells as well as the epithelial-like A549 and U2OS osteosarcoma cells exhibit diverse frequencies of lamina blebs after constricted migration (through 3 µm pores but not through 8 µm pores), all show a constriction-induced increase in cytoplasmic KU80 (versus nucleoplasmic) as well as an increase in γH2AX foci indicating an excess in DNA damage (Fig. 2A-iii). Cell densities were also always lowest after constricted migration, which has implications for cell cycle phase of the epithelial-like A549 and U2OS cells.

**Figure 2.**
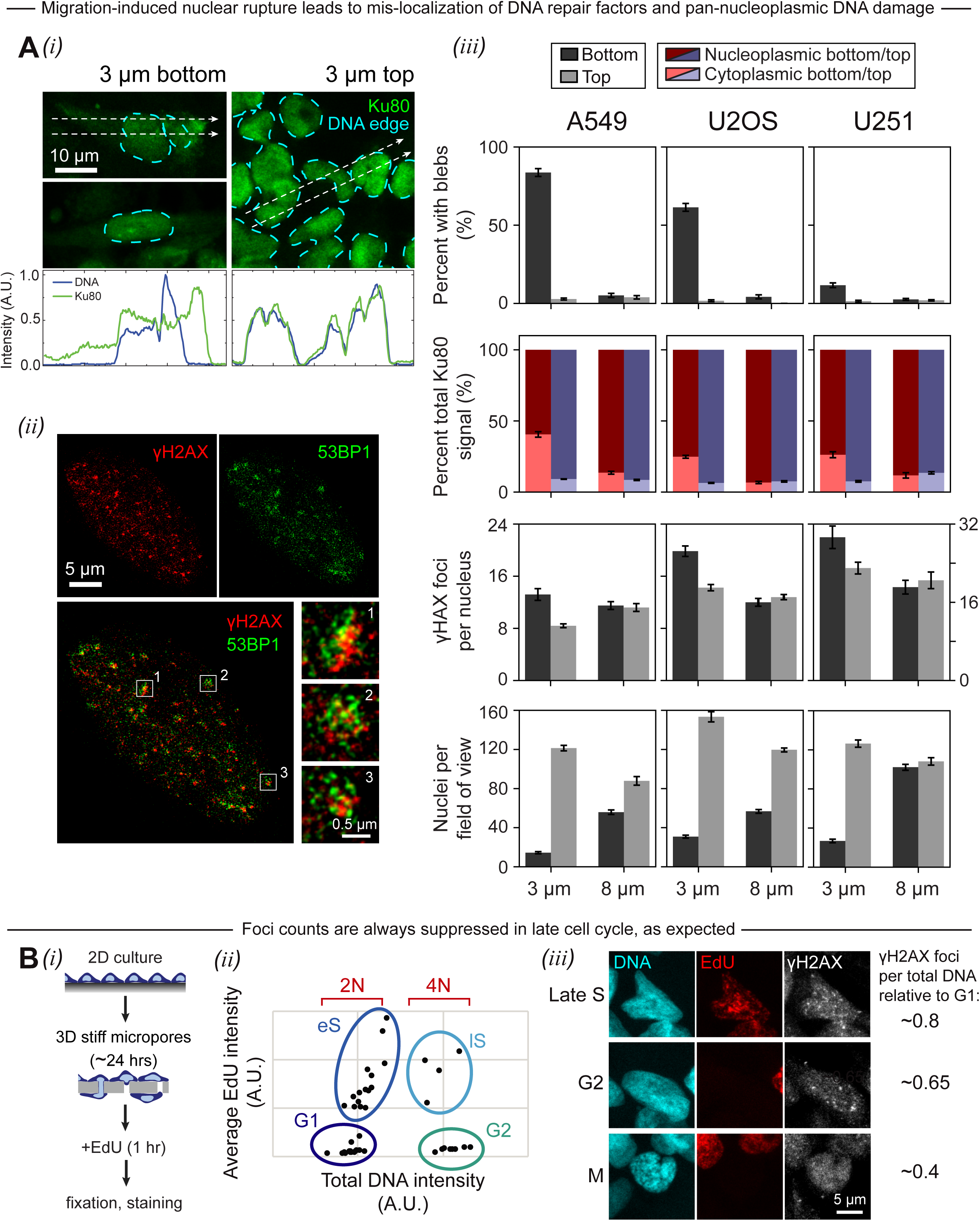
After 3 μm pore migration, three different cancer cell lines exhibit varying degrees of nuclear envelope rupture, leading consistently to repair factor mis-localization and pan-nucleoplasmic DNA double-strand breaks, with an expected decrease in foci per DNA in later phases of the cell cycle. (A) (i) Mobile nuclear proteins, including the DNA repair factor Ku80, are often observed outside the boundary of the DNA edge in cells that have migrated through 3 μm pores (‘3 μm bottom’). Such mis-localization, indicative of nuclear rupture, is rarely seen among unmigrated cells (‘3 μm top’). (ii) Super-resolution images of a representative formaldehyde-fixed, immuno-stained U251 cell show pan-nucleoplasmic 53BP1 foci that appear to mostly overlap with γH2AX foci. However, higher magnification insets reveal heterogeneity within individual foci. (iii) Three cancer cell lines—A549, U2OS, and U251 cells—exhibit diverse frequencies of lamina rupture following 3 μm pore migration (as indicated by the percent of migrated cells with blebs), but all exhibit a constriction-induced increase in the proportion of cytoplasmic (versus nucleoplasmic) endogenous KU80 as well as an increase in γH2AX foci. (B) (i) Schematic illustrating the EdU cell proliferation assay that was used to assess the impact of 3 μm pore migration on cell cycle progression. Immediately following a 24-hour migration period, EdU was added to the T ranswell membrane (or 2D culture) for 1 hour before the usual fixation and staining procedure. EdU-labeled cells were stained by a simple ‘click’ chemistry reaction that is perfectly compatible with immunohistochemical staining of other antigens. (ii) As shown in this representative plot, the EdU and DNA intensity of individual cells—measured by immunofluorescence microscopy—can be used to classify the cells as 2N or 4N and, further, as G1, early S (eS), late S (IS), or G2. (iii) As expected, cells in the later phases of the cell cycle (IS, G2, M) exhibit fewer γH2AX foci per total DNA content than cells in the earlier phases (G1, eS). Images show U251 cells on the top of an 8 μm pore Transwell membrane. Across all experimental conditions, U251 cells in late S have an average of 0.8* as many γH2AX foci per DNA as U251 cells in G1; cells in G2 have 0.65* as many; and mitotic cells have 0.4* as many. The trends are similar for A549 and U2OS cells.

Cell cycle analyses on the various pore migrated cells were done by quantitative fluorescence imaging of DNA content and DNA incorporation of EdU, as a measure of replication (Fig. 2B-i). Conventional designations for non-replicated genomes (2N) and fully replicated genomes (4N) are used for simplicity despite the aneuploid nature typical of cancer genomes (Fig. 2B-ii). As expected, the number of γH2AX foci per total DNA content decreases in the later cell cycle phases—late S (IS), G2, M—compared to G1 (Fig. 2B-iii). This is consistent with cell cycle checkpoints for DNA damage. However, it was also noticed that constricted migration also tended to suppress the proportion of cells in late phases of cell cycle (Fig. 3A-i).Cell cycle analyses on the various pore migrated cells were done by quantitative fluorescence imaging of DNA content and DNA incorporation of EdU, as a measure of replication (Fig. 2B-i). Conventional designations for non-replicated genomes (2N) and fully replicated genomes (4N) are used for simplicity despite the aneuploid nature typical of cancer genomes (Fig. 2B-ii). As expected, the number of γH2AX foci per total DNA content decreases in the later cell cycle phases—late S (IS), G2, M—compared to G1 (Fig. 2B-iii). This is consistent with cell cycle checkpoints for DNA damage. However, it was also noticed that constricted migration also tended to suppress the proportion of cells in late phases of cell cycle (Fig. 3A-i).

**Figure 3.**
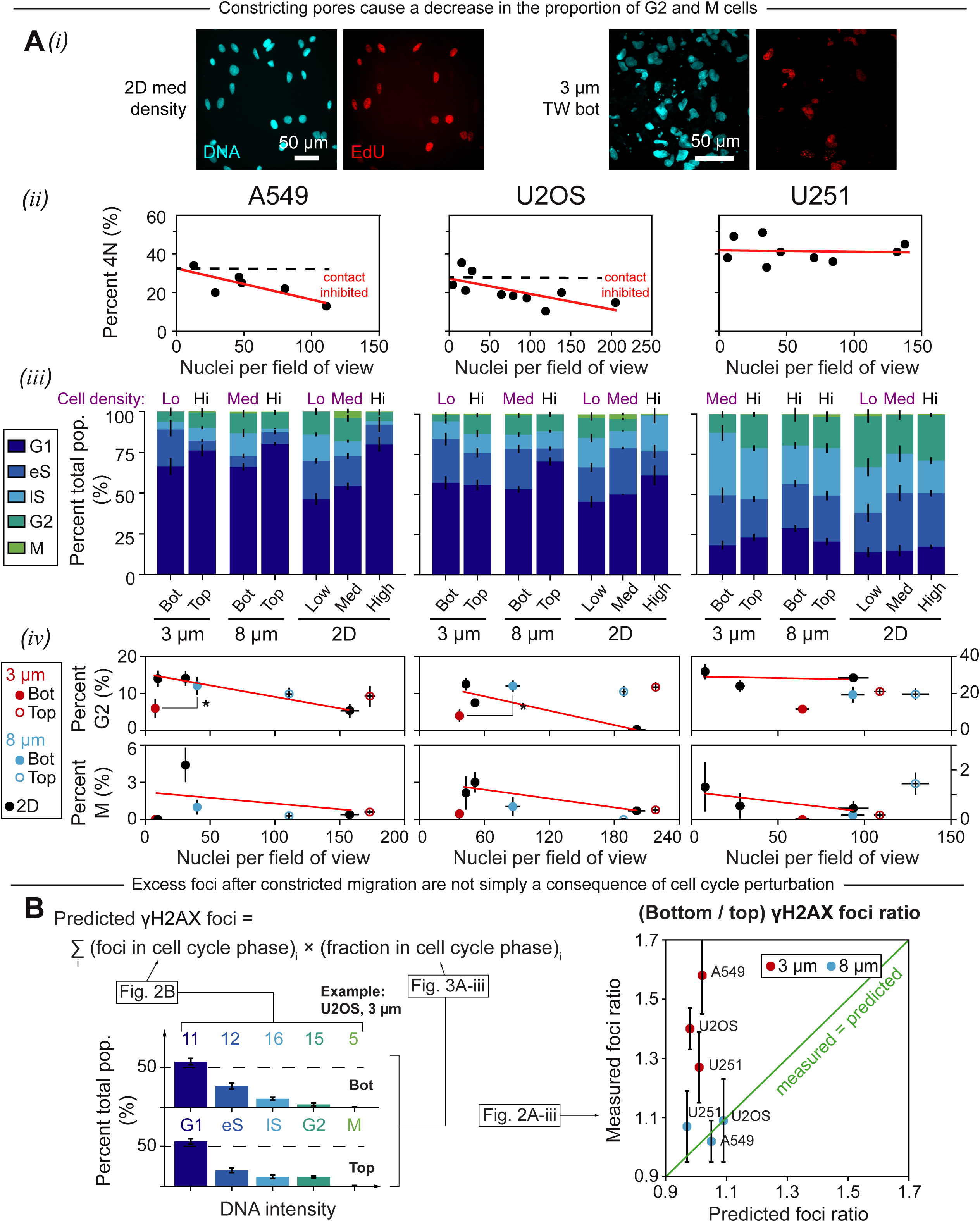
Constricted migration causes a perturbation to cell cycle, which does not explain away the constriction-induced increase in DNA damage. (A) A549, U251, and U2OS cells were seeded at three different densities (low, medium, high) on 2D plastic and were also migrated through 3 μm and 8 μm pore Transwell membranes. In each case, cell cycle analysis was performed using total DNA content combined with EdU incorporation, as described previously in Fig. 2 and as depicted here in panel (i). (ii) Epithelial-like A549 and U2OS cells seem to experience stronger contact inhibition than mesenchymal-like U251 cells. (iii-iv) Migration causes a decrease in the proportion of G2 and M cells for every cancer cell line studied, whether or not the cell line exhibits contact inhibition. Migration through the larger pores is consistent with cell cycle progression, with more cells in G2 phase compared to the small pores. (B) For each cell type, the average number of γH2AX foci was calculated for every phase of the cell cycle (G1, eS, lS, G2, M) across a number of experimental conditions. Based on these values, along with the cell cycle distributions from Fig. 3A-iii, the “predicted” number of γH2AX foci per nucleus was calculated for 3 μM and 8 μM top and bottom. Then, for both 3 μM and 8 μM, the measured (bottom/top) γH2AX foci ratio was plotted against the predicted foci ratio. Across all cell types, the 3 μM measured ratio is much larger than the predicted ratio, meaning that excess foci after constriction cannot be explained away as a consequence of perturbations to the cell cycle distribution.

A549 and U2OS cells are contact-inhibited in 2D cultures whereas U251 cells are not (Fig. 3A-ii), and so migration from high density (Top) to low density (Bottom) conditions after constricted migration should increase proliferation – at least for the epithelial-like cells. However, the proportions of G2 and M cells were suppressed for all lines studied, regardless of whether or not the cell line exhibits contact inhibition (Fig. 3A-iii,iv). Migration through the larger pores proved consistent with cell cycle progression, with more cells in G2 phase compared to the small pores.

Since different cell cycle phases have different numbers of γH2AX foci, as mentioned previously (Fig. 2B-iii), migration-induced changes in the cell cycle distribution should affect the average number of foci per nucleus. However, the predicted effect is small compared compared to the measured increase in γH2AX foci after 3 μm pore migration, meaning that excess foci after constriction is not merely a consequence of cell cycle perturbation (Fig. 3B). Moreover, the decreased number of cells in the cell cycle phases with the highest average foci counts (IS and G2) underscores the accumulation of damage in these phases or earlier.

To control cell cycle prior to migration, U2OS cells were plated at high density for 48 h to bias these contact-inhibited cells toward G1/eS. In 2D sparse cultures, the %-age of cells that are 4N decays in ~8 h toward the contact-inhibited state. When re-plated at high density above large pores, the cells migrate through to a low-density state and re-enter cell cycle and increase the %-age 4N, consistent with release from contact inhibition on Top (Fig. 4). However, constricting pores effectively repress cell cycle, with far fewer 4N cells, which is consistent with repression of G2/M phases (Fig. 3A-iii, iv).

**Figure 4.**
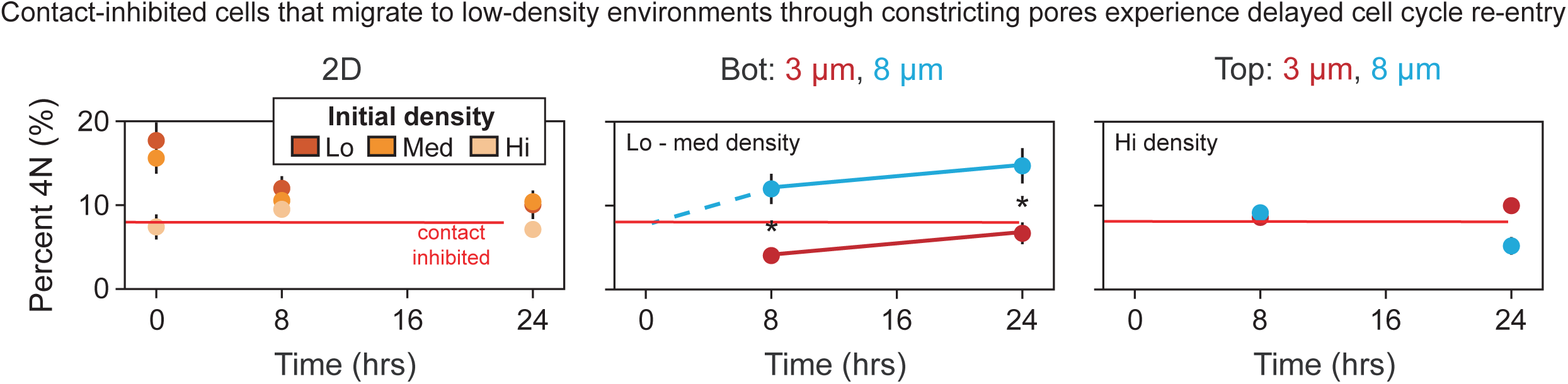
Starting with contact-inhibited cells, migration through large pores into sparse microenvironments releases contact inhibition and enables cell cycle re-entry, whereas constricted migration represses entry into late S/G2/M despite low density. U2OS cells were plated at high density for 48 hours to loosely synchronize them in G1/eS phase. After 48 hours, the cells were trypsinized; a portion of them were seeded on 3 and 8 μM TW membranes, and then allowed to migrate for 8 or 24 hours. Another portion of the cells were seeded at three different densities (high, med, low) in 2D, and then incubated for either 0, 8, or 24 hours. Migration through 8 μM pores causes an increase in the 4N population as cells move from high density on top of the Transwell membrane (right-most plot, contact-inhibited) to low density on the bottom (middle plot). Migration through 3 μM pores, despite also alleviating contact inhibition, slows entry into 4N over at least 24 hours.

To begin to test whether the excess DNA damage observed during constricted migration might relate to replication stress, siRNA was used to partially knock down one of the sources of DNA breaks in cells, the topoisomerase TOP2A. If TOP2A were indeed responsible for the excess foci, then siTOP2A cells would be expected to show a smaller increase in γH2AX foci after 3 μm pore migration than their wild-type counterparts (siCtrl or non-treated). However, partial depletion of TOP2A shows only a slight decrease in γH2AX foci and does not strongly affect cell cycle or the excess DNA damage after migration (Fig. 5).

**Figure 5.**
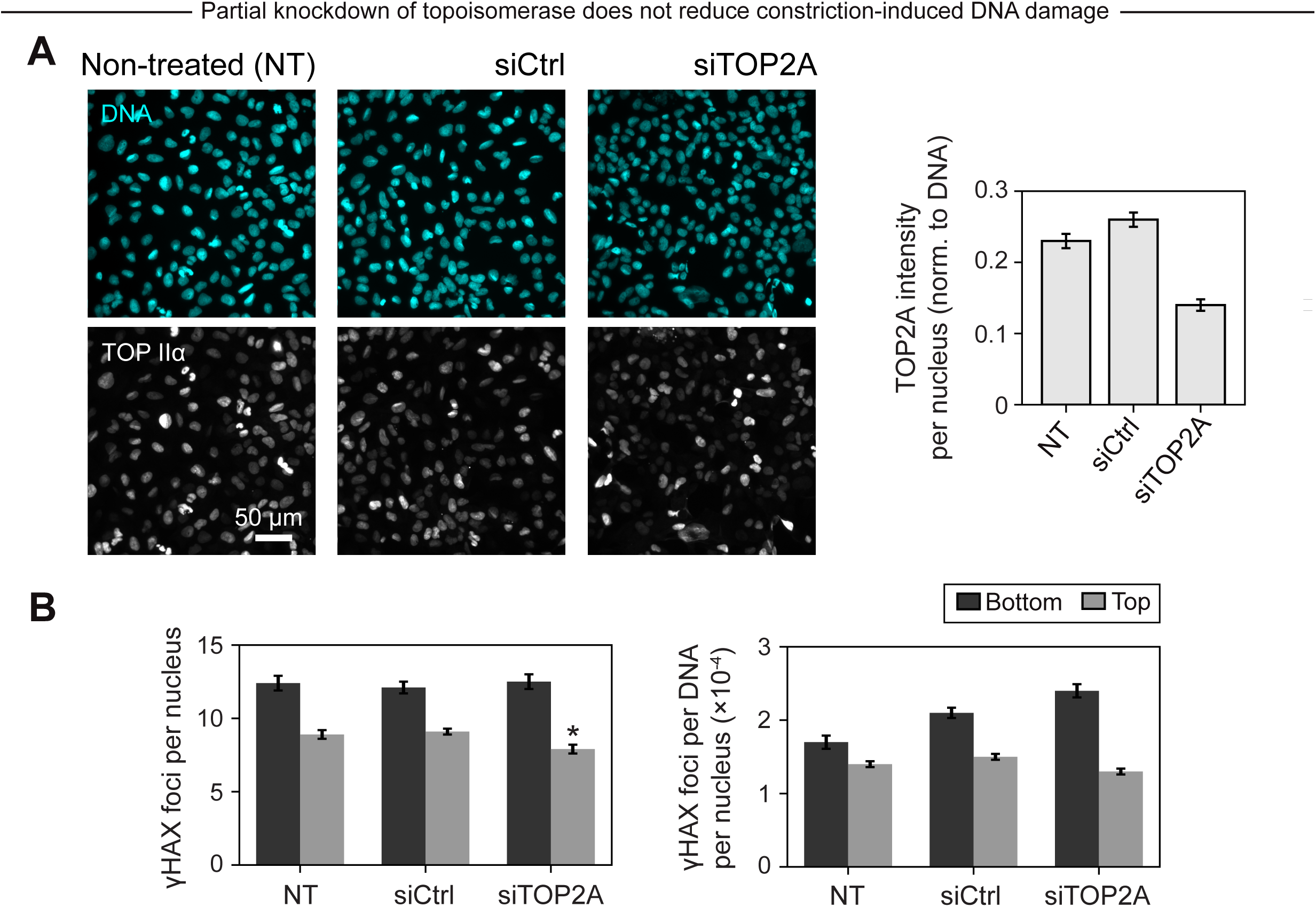
Partial knockdown of TOP2A does not eliminate DNA damage due to constricted migration. (A) siRNA was used to partially knock down the topoisomerase TOP2A. Knockdown efficiency was determined by immunofluorescence microscopy. (B) If TOP2A were responsible for the excess DNA double-strand breaks observed after after 3 μm pore migration, then siTOP2A cells would be expected to show a smaller increase in γH2AX foci after migration than their wild-type (non-treated or siCtrl) counterparts. However, partial depletion of topoisomerase does not strongly affect cell cycle or the excess DNA damage, consistent with weak dependencies on replication stress.

## Discussion

Small pores (3 μm) cause nuclear rupture unlike large pores (8 μm) (Fig.1,2A), and so it is possible that small pores mis-localize key regulators of cell cycle. For example, DNA repair factors are needed to help cells minimize DNA damage to pass through DNA checkpoints before mitosis (Fig.2B), and such factors clearly mis-localize in migration through small pores but not large pores (Fig.2A). Mis-localization of repair factors provides a mechanism for the increased DNA damage with small pores but not large pores even when corrected for cell cycle phase (Fig.3B). However, small pores might also select for G1/eS cells [15], in which case the effects of contact inhibition need to be carefully considered. Cells at high density on one side of a porous barrier will be enriched in G1 (or 2N), and such cells should re-enter cell cycle if separated out.

For large pores of 8 μm, there is not any obvious selection for G1/eS cells. After 24 h of migration, %G2 on ‘Bot’ is very similar to results expected from a low 2D density regardless of contact-inhibited cell type (Fig.3A-iv). For contact-inhibited cells, the %4N on ‘Bot’ after 8 h and 24 h is well above the contact-inhibited limit and thus consistent with cell cycle re-entry at low to medium density (Fig.4).

For small pores of 3 μm, G1/eS cells are somehow enriched. By 24 hours, %G2 on ‘Bot’ is well below results expected from 2D density independent of whether the cell type is contact-inhibited (Fig.3A-iv). For contact-inhibited cells, the %4N on ‘Bot’ after 8 h and 24 h are both well below the results for large pores (Fig.4), and despite the low densities, the 8-hour result is well below the contact-inhibited limit compared to %4N at 24 hours.

Small pores could select for G1/eS cells that then re-enter cell cycle or they could delay progression until DNA repair permits passage through checkpoints. Further experiments are needed to distinguish such possibilities, but it is clear that DNA damage after constricted migration is not an artifact of cell cycle perturbation – which certainly occurs.

## Materials and methods

### Cell culture

U2OS human osteosarcoma and U251 human glioblastoma cells were cultured in DMEM high glucose media (Gibco, Life Technologies), supplemented with 10% FBS and 1% penicillin/streptomycin (MilliporeSigma). A549 human lung cancer epithelial cells were cultured in supplemented Ham’s F12 media (Gibco, Life Technologies). A549 cells with endogenously GFP-tagged CTNNB1 and RFP-tagged LMNB1 were purchased from MilliporeSigma.

### Transfection

GFP-53BP1 was a gift from Dr. Roger Greenberg of the University of Pennsylvania [16]. siTOP2A (30 nM, Santa Cruz), ON-TARGETplus Non-targeting Control Pool (30 nM, Dharmacon), or GFP-53BP1 (0.5 ng/mL) were transfected by Lipofectamine 2000 (Invitrogen, Life Technologies) for 3 days (siRNA) or 24 hours (GFP) in corresponding media, supplemented with 10% FBS. Knockdown efficiency was determined by immunofluorescence microscopy.

### Transwell migration

Cells were plated on top of a Transwell membrane (Corning) at a density of 300,000 cells/cm^2^ and allowed to migrate to the bottom over the course of 24 hours.

### Immunostaining and imaging

Cells were fixed in 4% formaldehyde (MilliporeSigma) for 15 minutes before undergoing 10-minute permeabilization by 0.25% Triton-X (MilliporeSigma), 30-minute blocking by 5% BSA (MilliporeSigma), and overnight incubation in primary antibodies. The antibodies used include lamin-A/C (Santa Cruz and Cell Signaling), Lamin-B (Santa Cruz), γH2AX (MilliporeSigma), 53BP1 (Abcam), and topoisomerase IIα (Santa Cruz). The cells were then incubated in secondary antibodies (Thermo Fisher) for 1.5 hours, and their nuclei were stained with 8 μM Hoechst 33342 (Thermo Fisher) for 15 minutes. Finally, the cells were mounted with Prolong Gold antifade reagent (Invitrogen, Life Technologies). Epifluorescence imaging was done using an Olympus IX71—with a 40x/0.6 NA objective—and a digital EMCCD camera (Cascade, Photometrics). Confocal imaging was done on a Leica TCS SP8 system, equipped with either a 63x/1.4 NA oil-immersion or a 40x/1.2 NA water-immersion objective. Super-resolution images were taken using a Leica TCS SP8 STED 3X system with a 100x/1.4 NA oil-immersion objective. ImageJ [17] was used to quantify the resulting images.

### EdU labeling and staining

EdU (10 μM, Abcam) was added to 2D culture or Transwell membrane 1 hour before fixation and permeabilization. After permeabilization, samples were stained with 100 mM Tris (pH 8.5) (MilliporeSigma), 1 mM CuSO4 (MilliporeSigma), 100 μM Cy5 azide dye (Cyandye), and 100 mM ascorbic acid (MilliporeSigma) for 30 min at room temperature. Samples were thoroughly washed, and then underwent immunostaining as described above.

### Micropipette aspiration

In preparation for aspiration, cells were: (1) detached using 0.05% Trypsin-EDTA (Gibco, Life Technologies); (2) incubated in 0.2 μg/mL latrunculin-A (MilliporeSigma) and 8μM Hoechst 33342 (Thermo Fisher) for 30 minutes at 37°, as described previously [18]; and (3) re-suspended in PBS with 1% BSA and 0.2 μg/mL latrunculin-A. During aspiration, epifluorescence imaging was done using a Nikon TE300—with a 60x/1.25 NA oil-immersion objective—and a digital EMCCD camera (Cascade, Photometrics). The resulting images were quantified in ImageJ [17].

